# Unc-51-like Kinase 1 (ULK1) Regulates Bacterial Ubiquitylation and p62 Recruitment during Xenophagic Clearance of *Listeria monocyt*ogenes

**DOI:** 10.1101/2024.10.23.619899

**Authors:** MS Siqueira, TSM Farias, RM Ribeiro, JS Ribeiro, TD Pereira, LAM Carneiro, LH Travassos

**Affiliations:** Laboratory of Immunoreceptors and Signaling, Institute of Biophysics Carlos Chagas Filho, Federal University of Rio de Janeiro, Brazil; Laboratory of Innate Immunity and Inflammation, Department of Immunology, Institute of Microbiology Paulo de Góes, Federal University of Rio de Janeiro, Brazil

## Abstract

Autophagy is an essential cellular homeostatic process, that also serves as an innate immune mechanism against intracellular bacterial pathogens, through a highly selective form of autophagy known as xenophagy. Despite advances in understanding how bacteria are targeted for autophagic degradation, the specific regulatory mechanisms that drive the initial steps and ensure bacterial selection remain incompletely defined. Our study uncovers a pivotal role for Unc-51-like kinase 1 (ULK1) in the xenophagic clearance of the intracellular bacterial pathogen *Listeria monocytogenes*. We observed that ULK1 is essential for the efficient ubiquitylation of bacteria and subsequent recruitment of the autophagic adaptor protein p62 to the bacterial surface. Furthermore, we show that the impact of ULK1 deficiency in these early events - reduction in bacterial ubiquitylation followed by impaired p62 targeting – later result in diminished formation of bacteria-targeted autophagosomes. Notably, phosphorylation of p62 at the S409 residue, which is known to enhance its affinity for ubiquitin, is necessary for its recruitment and effective bacterial clearance, highlighting the regulatory role of ULK1 in this process. These findings unveil a previously unrecognized function of ULK1 in modulating early xenophagy steps, contributing to the autophagic control of intracellular pathogens. Our findings offer new perspectives into the manipulation of ULK1activity for therapeutic interventions against infectious diseases.

**IMPORTANCE:** Autophagy is a vital process in eukaryotic cells that enables them to digest intracellular components, helping them respond to various stresses, including starvation, the accumulation of dysfunctional organelles, and infections. While the autophagic flux has been extensively studied over the past few decades, some key mechanisms remain poorly understood. In our research, we aimed to clarify one such mechanism: the way the autophagic machinery specifically targets intracellular bacteria. We identified a novel role for the protein ULK1 in this process, demonstrating that ULK1 is essential for marking bacteria within the cell and recruiting the protein p62. This marking and recruitment of p62 are critical steps for effective bacterial clearance. This underscores the pivotal role of ULK1 in initiating the cellular defense against bacterial infections. Our findings could pave the way for new therapeutic strategies aimed at enhancing the body’s capacity to combat bacterial infections.

## INTRODUCTION

Autophagy is a cellular process highly conserved in eukaryotes that maintains homeostasis at cellular and systemic levels. During its activation, portions of the cytosol are sequestered into double-membrane vesicles called autophagosomes that can either fuse with late endosomes to form amphisomes or directly with lysosomes, resulting in the degradation of its content (1). Although firstly thought of as a non-selective bulk degradation process responsible for maintaining energy levels during nutrient deprivation, in the last two decades, evidence accumulated that autophagy can be a highly selective process in which a variety of adaptor proteins recruit the autophagy machinery to specific cargos (2).

Studies from the early 2000s demonstrated that autophagy could target several intracellular bacterial pathogens, such as group A *Streptococcus* (3), *Mycobacteria tuberculosis* (4), and *Shigella flexneri* (5), in a specific form of autophagy, called xenophagy. These studies were instrumental in establishing xenophagy as a vital effector mechanism of the innate immune defense arsenal.

Recognition of bacteria or bacterial products by surface (6) or cytosolic pattern recognition receptors (PRRs) (7, 8) have been implicated in the initiation of xenophagy. Still, they alone cannot explain how the formation of new autophagosomes is directed to specifically engulf bacteria in the cytosol. Much effort has been put into understanding the initial steps of xenophagy, mainly how bacteria are targeted as cargo for autophagy. Currently, it is known that the activity of different ubiquitin ligases such as leucine-rich repeat and sterile-motif–containing 1 (LRSAM1) (9), Parkin (10), SMAD-specific E3 ubiquitin protein ligase 1 (Smurf1) (11), Nedd4-1 (12) results in the decoration of intracellular bacteria with ubiquitin. This is a crucial event to target cytosolic bacteria to autophagosomes because it allows autophagic adaptors, such as p62 (13), nuclear dot protein 52 kDa (NDP52) (14), and optineurin (15), which harbor both ubiquitin-binding domains and LC3-interacting region (LIR) motifs in their structure, to act as bridges between the ubiquitylated bacteria and phagophores (16). Despite all these advances in the knowledge regarding the involvement of ubiquitin-ligases and autophagic adaptors, much is to be learned concerning the players that regulate this critical step for xenophagy.

Unc-51-like kinase 1 (ULK1) is a serine-threonine kinase that is part of the ULK1 complex, consisting of ULK1, the non-catalytic focal adhesion kinase family interacting protein of 200 kD (FIP200), ATG13 and ATG101 (17). This core autophagy initiation complex is present on phagophore membranes and, in concert with class III phosphatidylinositol 3-kinase complex (Beclin 1, ATG14L, Vps34, and Vps15), regulates the early steps of autophagosome formation (17, 18). Phagophore expansion into autophagosome requires additional proteins such as WD-repeat protein Interacting with PhosphoInositides (WIPI), ATG9, ATG2, and two ubiquitin-like systems: the ATG12-ATG5 system, in which ATG12 is conjugated to ATG5 to form a complex consisting of ATG12, ATG5, and ATG16L1. This complex is necessary for the conjugation of phosphatidylethanolamine to the ATG8 system, a family of proteins present on the membrane of the nascent autophagosome (19).

Previous studies demonstrate that the ULK1 complex detects upstream signals from energy-sensing pathways, such as mechanistic Target of Rapamycin (mTOR) and AMP-activated protein kinase (AMPK), thus playing a central role in regulating intracellular energy levels (17). mTOR and AMPK are master regulators of nutrient/energy levels and have been demonstrated to regulate ULK1 activity through phosphorylation at different sites (20–23). Although the essential role of ULK1 in the initiation of starvation-induced autophagy is undisputed, its participation in bacteria-targeted xenophagy is far from being elucidated. Here, we investigated the role of ULK1 in xenophagy, more specifically in how it participates in the initial events of bacterial ubiquitylation and p62 recruitment to bacterial surface that will result in bacterial clearance by autophagy. For this purpose, we used *Listeria monocytogenes,* an intracellular bacterial pathogen that escapes from phagosomes to replicate in the cytosol of host cells. This bacteria represents an excellent experimental model as it has long been reported to trigger ubiquitylation and p62 recruitment to its surface, leading to the formation of autophagosomes around bacteria (24). We demonstrate that infection of mouse embryonic fibroblasts (MEFs) with *L. monocytogenes* leads to the recruitment of ubiquitin and p62 to the bacterial surface, forming LC3^+^ compartments colocalizing with bacteria. Moreover, we found that this process depends on the escape to the cytosol as phagosome-confined *L. monocytogenes* do not recruit xenophagy markers nor colocalize to autophagosomes. More importantly, we show that, in ULK1-deficient MEFs, bacteria is not efficiently ubiquitylated, and p62 is not targeted to the bacterial surface throughout infection with *L. monocytogenes*. In addition, we demonstrate that the phosphorylation of p62 at the S409 residue, known to be dependent on ULK1 (25) and to increase its affinity to ubiquitin, is required to recruit p62 to the bacterial surface. Finally, we demonstrate that ULK1 deletion or the S409A substitution that impairs p62 phosphorylation resulted in significantly increased numbers of intracellular bacteria. Together, our results point to a yet unrecognized role of ULK1 in controlling bacterial ubiquitylation and p62 recruitment to the surface of cytosolic bacteria, which contributes to bacterial control

## RESULTS

### Cytosolic *L. monocytogenes* triggers xenophagy

It has been described that *L. monocytogenes* and other intracellular bacteria interact with the autophagic machinery in multiple ways (24–26). These bacteria have evolved mechanisms that allow them to manipulate this host response and, in some cases, even take advantage of the autophagy response. To quantify *L. monocytogenes* targeting and kinetics of autophagy response in our model, we infected wild-type MEFs to evaluate the recruitment of the autophagy marker LC3 to the bacterial surface. Immunofluorescence experiments demonstrated the association of LC3 with the wild-type strain of *L. monocytogenes*, indicating that xenophagy was triggered at 2h (**Figure 1A and D**). To precisely determine the dynamics of LC3 recruitment to the bacterial surface, MEFs were infected for 1, 2, 4, or 8h, and the number of cytosolic bacteria associated with LC3 was determined by immunofluorescence. As shown in **Figure 1B**, *L. monocytogenes* association with LC3 was maximal at 4h post-infection (∼10% of total bacteria). Xenophagy can target different intracellular bacterial species that survive in the cytosol (7) or damaged vacuoles (27). To determine in which intracellular compartment *L. monocytogenes* was targeted by xenophagy, we used a mutant of *L. monocytogenes* that lacks listeriolysin O (LLO), a cholesterol-dependent pore-forming cytolysin (encoded by *hly* gene) that is unable to escape from phagosomes and access the host cytosol, as previously demonstrated (28–30). We confirmed that bacteria retained in intact vacuoles do not initiate autophagy as Δ*hly L. monocytogenes* did not associate with LC3 (**Figure 1 C and E**).

**FIGURE 1:**
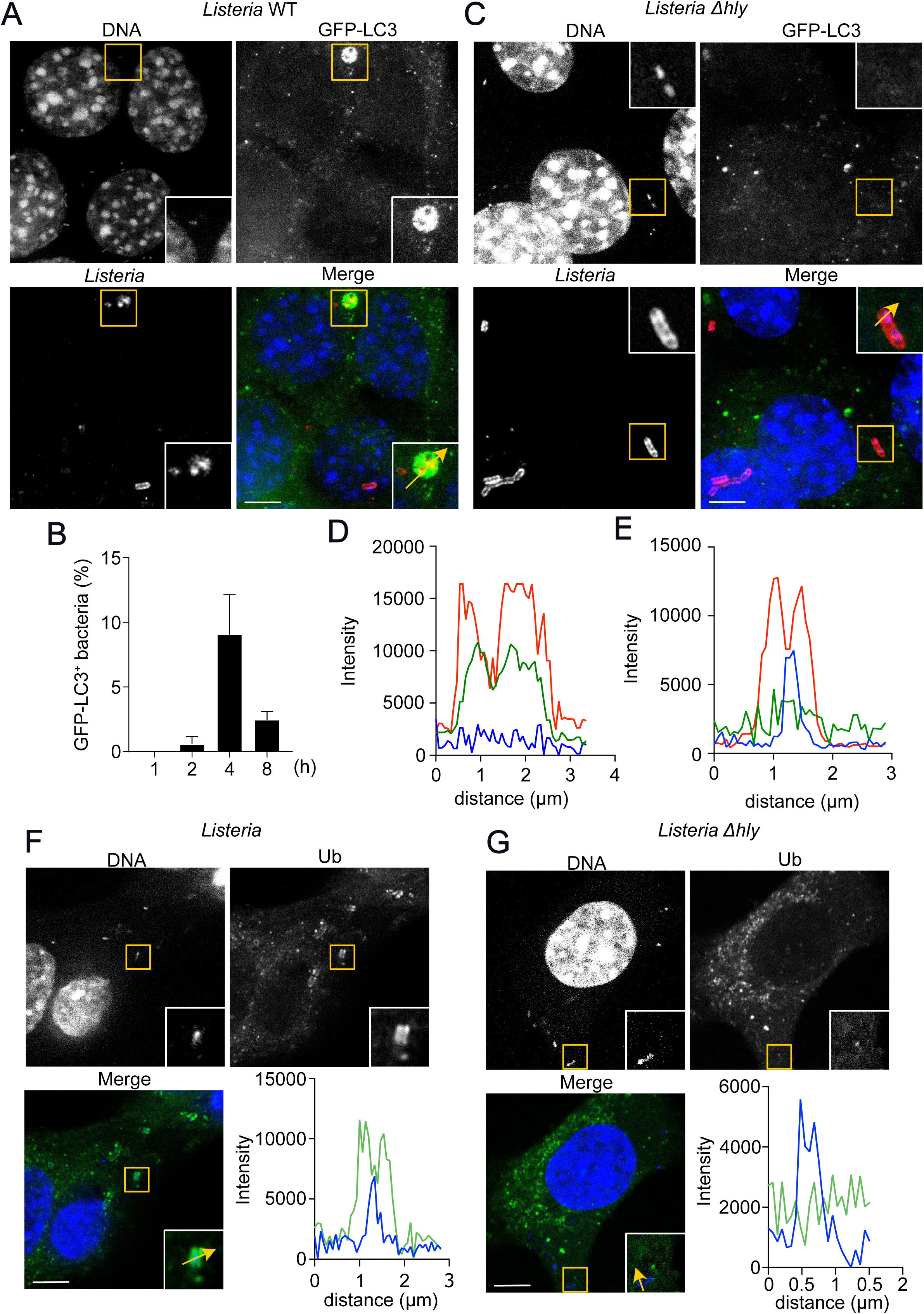
*L. monocytogenes*-targeted autophagy depends on listeriolysin O (LLO) expression. MEFs transduced with GFP-LC3 were infected with 10403S (WT) or 10403S Δ*hly* (Δ*hly*) *L. monocytogenes* and stained at 4h post-infection. (a) Representative confocal images of MEFs GFP-LC3 infected with WT or Δ*hly* (c) and stained for *Listeria* (red) and DNA (blue), N=2. (b) Quantification of the number of intracellular *L. monocytogenes* colocalizing with GFP-LC3 at 1, 2, 4, and 8 h, N=2. (d) and (e) Plot-profiles of fluorescence intensity along the yellow arrow traced in the insets in (a) (WT) and (b) (Δ*hly*); GFP-LC3 (green line), *Listeria* (red line), and DNA (blue line). Representative confocal images and plot profiles of fluorescence intensity along the yellow arrow traced in the insets of MEFs infected with WT (f) or Δ*hly* (g) for 2h and stained for ubiquitin (green) and DNA (blue), N=3. Values are means ± SEM. Scale bars are 10μM.

### Ubiquitin and p62 are recruited to *L. monocytogenes* surface

Since the description of xenophagy as a cell-autonomous mechanism that deals with intracellular bacteria, it has become clear that early events of bacterial ubiquitylation are essential to allow the recruitment of autophagy adaptors that will direct the formation of the autophagosomes around bacteria. To understand these crucial events for *L. monocytogenes*-induced autophagy, we started by analyzing the dynamics of ubiquitin (Ub) recruitment during the infection. To this end, WT MEFs infected with either wild-type or Δ*hly L. monocytogenes* were stained for ubiquitin at different time points, and the number of ubiquitin-positive (Ub^+^) bacteria was quantified. Consistent with the findings described above, the LLO-deficient strain of *L. monocytogenes* was not stained with Ub, in sharp contrast to what was observed with the wild-type strain, which had ∼50% of the intracellular bacterial population Ub^+^ as early as 2h post-infection (**Figure 1F,G** and **Supp. Fig. 1A-C**). Once ubiquitylated, bacteria may be targeted to autophagy by several ubiquitin-binding adaptors. Among them, p62 emerges as the best characterized one and has been consistently reported to bind ubiquitin-coated bacteria in the cytosol (31). Thus, next, we analyzed the recruitment of p62 to the surface of Ub^+^ *L. monocytogenes*. For this purpose, we immunostained WT MEFs infected with wild-type *L. monocytogenes* for Ub and p62. Our analysis demonstrates that most of the bacterial population (∼65%) was Ub^+^p62^+^, in line with the notion that p62 is implicated in recognizing ubiquitylated cytosolic bacteria (**Figure 2A-D**).

**FIGURE 2:**
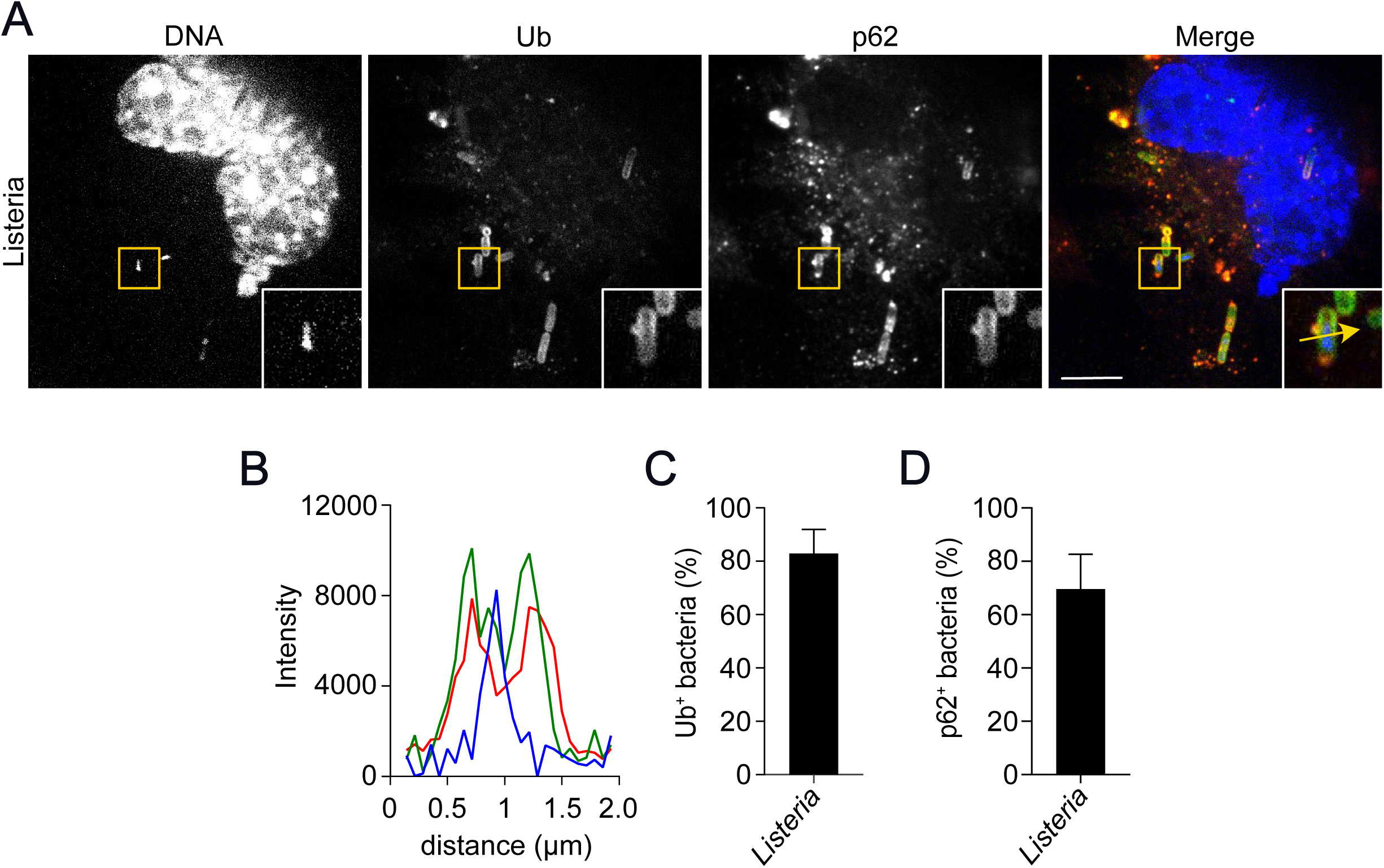
p62 is recruited to the surface of *L. monocytogenes* early during infection. (a) Representative confocal images of MEFs infected with WT *L. monocytogenes* for 2h and stained for ubiquitin (green), p62 (red), and DNA (blue), N=3. (b) Plot profiles of fluorescence intensity along the yellow arrow traced in the inset in (a); ubiquitin (green line), *Listeria* (red line), and DNA (blue line). (c) and (d) Quantification of the number of Ub^+^ (c) and p62^+^ (d) *L. monocytogenes*, N=3. Values are mean ± SEM. Scale bars are 10μM.

### *Listeria monocytogenes* induces delocalization of mTOR from lysosomes

Despite compelling evidence demonstrating that cytosolic bacteria are ubiquitylated and later targeted by different autophagic adapters, little is known concerning the factors controlling this critical step of xenophagy. Autophagy is a cellular stress response activated due to an intricated signaling network governed by two major complexes, mTOR and AMPK, that are crucial regulators of cellular protein synthesis and energy levels, respectively (20, 21). Downstream of them, several other signaling complexes integrate multiple inputs to induce or inhibit the formation of autophagosomes. One of these complexes is the ULK1-FIP200-ATG13 complex. Under nutrient-rich conditions, mTOR phosphorylates the serine-threonine kinase ULK1 to repress its activity, inhibiting autophagy induction (20, 21). Under amino acid starvation, however, this inhibitory phosphorylation is released, allowing ULK1 to initiate autophagosome formation. Since it has been demonstrated that infection with intracellular bacterial pathogens triggers amino acid starvation followed by mTOR inhibition (32), and this is known to activate ULK1 to initiate autophagosome formation, we hypothesized that ULK1 could connect the recruitment of ubiquitin and p62 to the surface of *L. monocytogenes* with autophagosome seeding. To test this hypothesis, we first asked if *L. monocytogenes* infection triggered amino acid deprivation. To this end, we investigated the delocalization of mTOR from lysosomes during *L. monocytogenes* infection, an event that has been shown to correlate with mTOR inhibition by *L. monocytogenes*-induced amino acid deprivation (32). Immunofluorescence analysis of mTOR and LAMP1 distribution in WT MEFs infected with *L. monocytogenes* revealed that mTOR localizes to lysosomes (∼56%) under nutrient-rich conditions, as observed in **Figure 3A**. However, *L. monocytogenes* infection for 1h resulted in a rapid displacement of mTOR from lysosomes, with only 16% of the infected cells displaying colocalization of mTOR and LAMP1. Interestingly, the absence of colocalization persisted throughout the infection, peaking at 4h when only ∼3% of the infected cells showed mTOR colocalizing with Lamp1 (**Figure 3B-F**). Our results confirm that *L. monocytogenes* infection reduces the colocalization of mTOR and lysosomes, likely due to amino acid deprivation as previously described (32).

**FIGURE 3:**
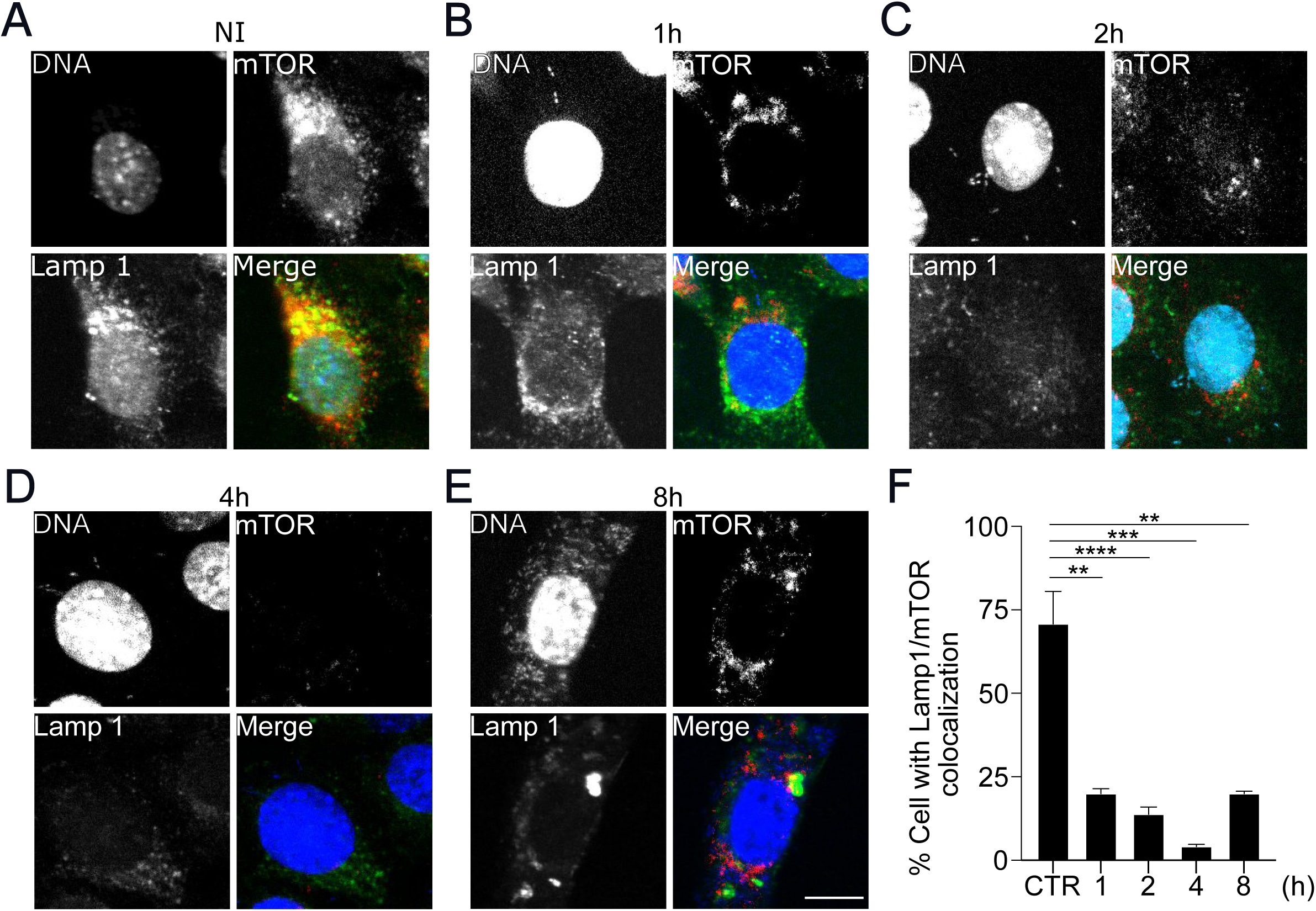
*L. monocytogenes* infection reduces the colocalization of Lamp1 and mTOR to induce amino acid starvation. (a) Representative confocal images of MEFs left uninfected and stained for Lamp1 (green), mTOR (red), and DNA (blue). (b-e) Representative confocal images of MEFs infected for 1h (b), 2h (c), 4h (d), and 8h (e) and stained as in (a). (f) Quantification of the number of cells displaying Lamp1/mTOR colocalization. Values are mean ± SEM. Scale bars are 10μM. **p<0.01, ***p0.001 (one-way ANOVA with Dunnett’s post-test over CTR), N=2

### ULK1 is essential for ubiquitylation and recruitment of p62 to *L. monocytogenes* surface

The reduction in colocalization between mTOR and Lamp1 during *L. monocytogenes* infection and the fact that mTOR inhibition leads to ULK1 activation and autophagosome formation led us to hypothesize that ULK1 could also play a role in the ubiquitylation and recruitment of p62 to *L. monocytogenes* surface. To test this, we infected WT MEFs and MEFs from ULK1-deficient mice with wild-type *L. monocytogenes* and analyzed the recruitment of these two proteins by immunofluorescence. As shown in **Figures 4A**, **B**, **D**, and **E** at 2h post-infection, ∼70% of the bacterial population in the cytosol of WT MEFs were coated with ubiquitin and ∼60% with p62. In contrast, we observed that only ∼27% and ∼31% of *L. monocytogenes* in the cytosol of ULK1-deficient MEFs were decorated with Ub and p62, respectively (**Fig. 4A, C, D**, and **E**). To investigate whether this reduction in bacterial ubiquitylation and p62 recruitment in ULK1-deficient MEFs was due to delayed kinetics, we infected wild-type and ULK1-deficient MEFs for 4h and 8h and observed similar results. At 4h post-infection, we found that ∼73% vs. ∼22% of bacteria were positive for ubiquitin and ∼57% vs. ∼13% positive for p62 in WT vs. ULK1-deficient MEFs, respectively. Similar results were observed at 8h post-infection (**Fig 4F-I** and **Supp. Fig. 2A** and **D**).

**FIGURE 4:**
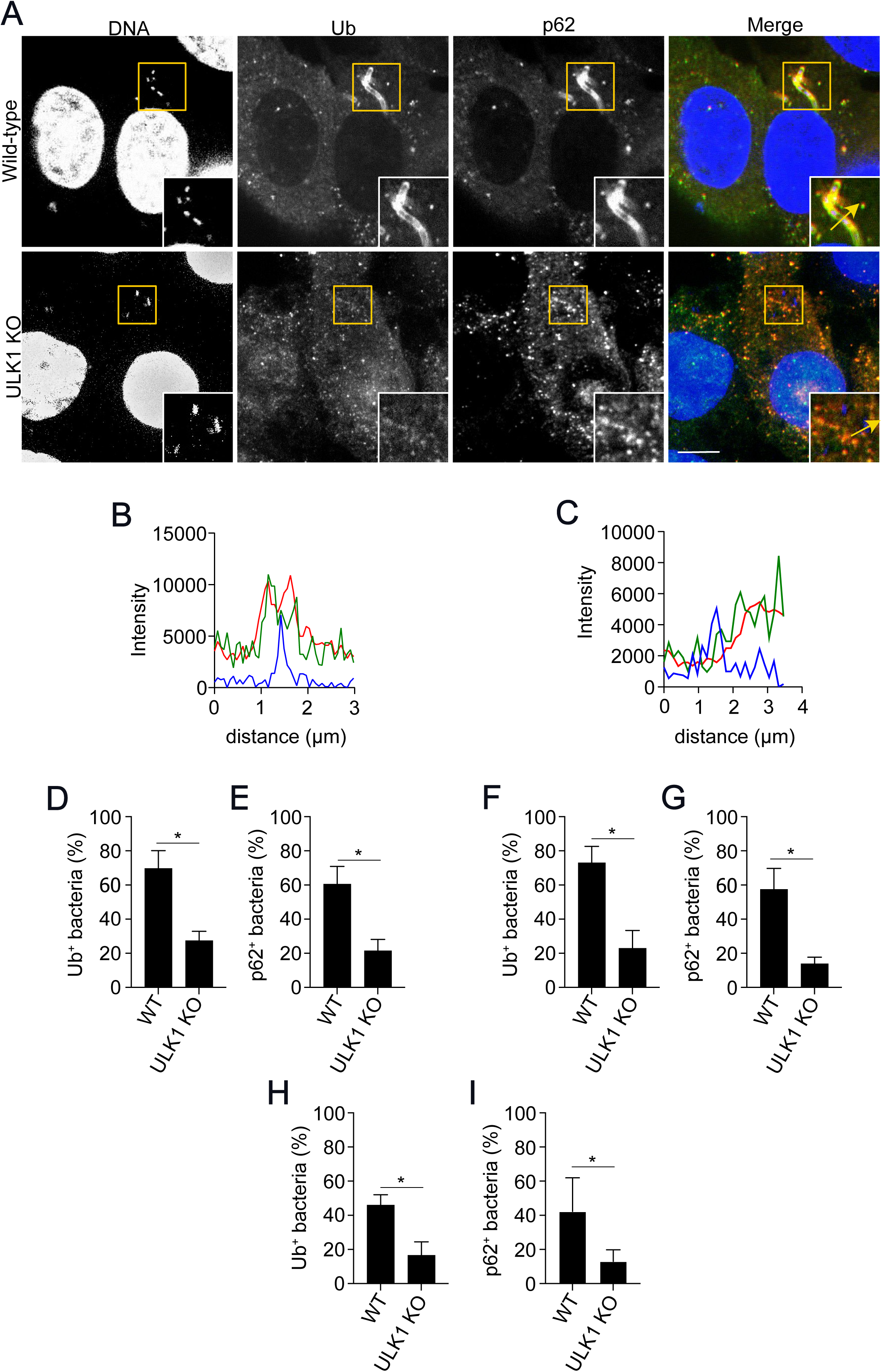
ULK1 is required for efficient recruitment of ubiquitin and p62 to *L. monocytogenes* surface. (a) Representative confocal images of WT and ULK1 KO MEFs infected with WT *L. monocytogenes* for 2h and stained for ubiquitin (green), p62 (red), and DNA (blue). N=5. (b) and (c) Plot profiles of fluorescence intensity in WT (b) and ULK1 KO MEFs (c) infected with *L. monocytogenes* along the yellow arrow traced in the inset in (a); ubiquitin (green line), *Listeria* (red line), and DNA (blue line). (d) and (e) Quantification of the number of WT *L. monocytogenes* positive for ubiquitin (d) and p62 (e) in WT and ULK1 KO MEFs infected for 2h. (f) and (g) Quantification of the number of Ub^+^ WT *L. monocytogenes* (f) and p62^+^ (g) in WT and ULK1 KO MEFs infected for 4h. (h) and (i) Quantification of the number of WT *L. monocytogenes* positive for ubiquitin (h) and p62 (i) in WT and ULK1 KO MEFs infected for 8h. Scale bars are 10μM. Values are mean ± SEM. *p<0.05, two-tailed unpaired t-test. N=3-5.

### ULK1 is not required for the ubiquitylation and recruitment of p62 to *L. monocytogenes* lacking ActA

Upon infection, *L. monocytogenes* mediates its escape from the entry vacuole by secreting LLO and two phospholipases C (PlcA and PlcB) (33, 34). Once free in the cytosol, *L. monocytogenes* masks its surface with ActA, a bacterial protein that promotes host actin polymerization that is used by the bacteria for its intracellular motility (35). Interestingly, this ActA mask has been shown to avoid Ub accumulation and p62 recruitment to bacterial surfaces (36, 37). Therefore, we aimed to understand if an ActA expression would affect the ULK1-dependent ubiquitylation and p62 recruitment to the *L. monocytogenes* surface. We used a Δ*actA* mutant to infect WT MEFs for different time points and monitor Ub and p62 coating of bacteria by immunofluorescence. The analysis of the colocalization of Ub and p62 with Δ*actA L.monocytogenes* revealed that virtually all bacteria were densely coated by both proteins throughout the infection (**Supp. Fig 3A-D**), confirming previous findings that the lack of ActA favors bacterial coating with Ub and p62. Next, we infected WT and ULK1 KO MEFs with the Δ*actA L. monocytogenes* mutant and analyzed the recruitment of Ub and p62. Immunofluorescence and plot profile analysis demonstrates that ULK1-deficiency does not impact the ubiquitylation and p62 recruitment to the surface of Δ*actA L. monocytogenes* (**Fig. 5A-C**). To confirm these findings, we quantified the number of Δ*actA* bacteria associated with Ub and p62. In sharp contrast to the results with wild-type bacteria (**Fig. 4**), ULK1 was not required for the ubiquitylation of Δ*actA L. monocytogenes* and p62 recruitment, as demonstrated by the similar recruitment of Ub and p62 in WT and ULK1 KO MEFs infected with Δ*actA L. monocytogenes* (**Fig. 5D-I** and **Supp. Fig. 4A** and **B**). Together with the results from previous sections, our data clearly demonstrate that ULK1 plays a crucial role in the early steps of xenophagy by controlling the ubiquitylation and recruitment of p62 to the surface of *L. monocytogenes* expressing ActA.

**FIGURE 5:**
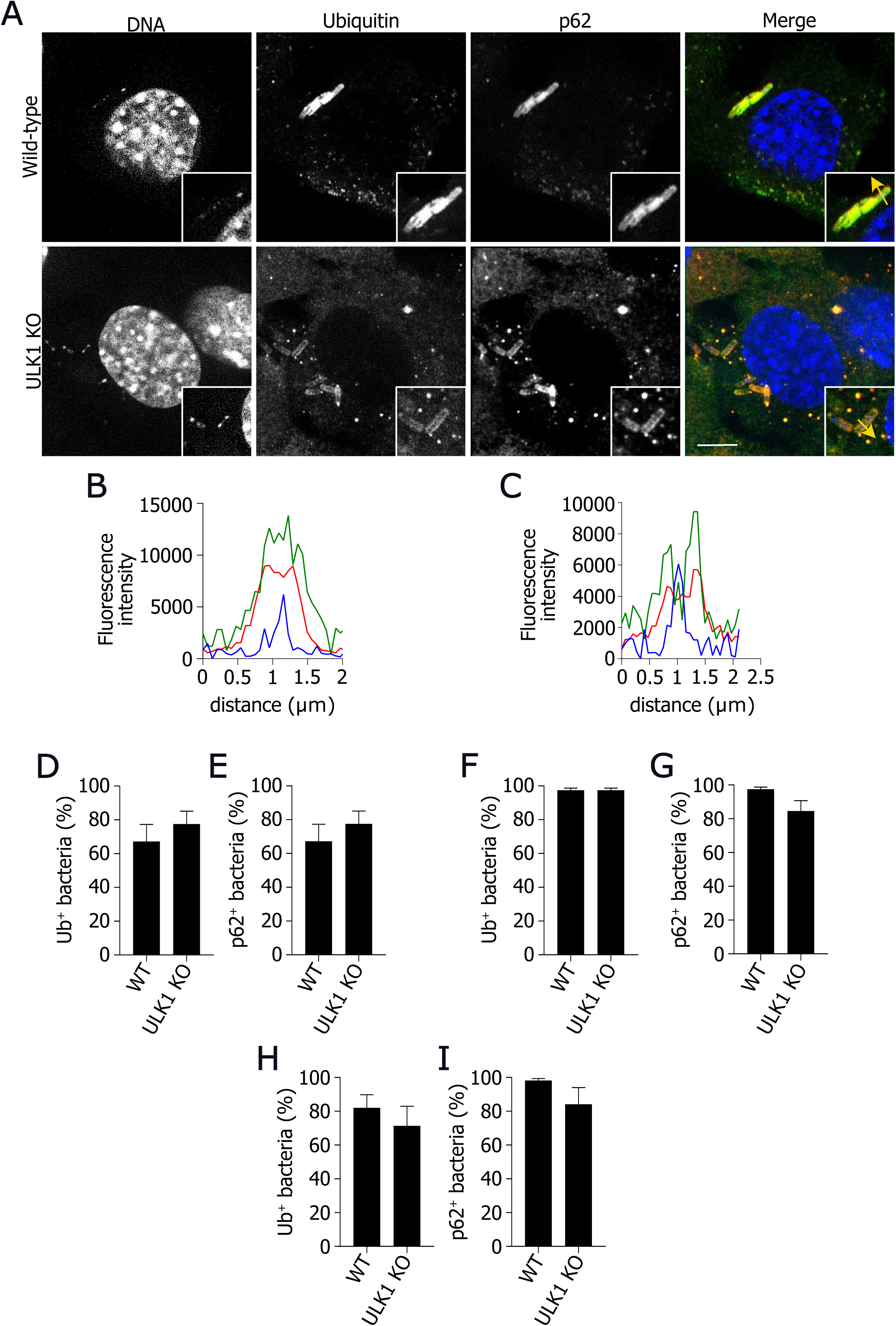
Ubiquitin and p62 recruitment to *L. monocytogenes* lacking actA do not require ULK1. (a) Representative confocal images of WT and ULK1 KO MEFs infected with WT Δ*actA L. monocytogenes* for 2h and stained for ubiquitin (green), p62 (red), and DNA (blue). N=5. (b) and (c) Plot profiles of fluorescence intensity in WT (b) and ULK1 KO MEFs (c) infected with *L. monocytogenes* along the yellow arrow traced in the inset in (a); Ubiquitin (green line), *Listeria* (red line), and DNA (blue line). (d) and (e) Quantification of the number of Δ*actA L. monocytogenes* positve for ubiquitin (d) and p62 (e) in WT and ULK1 KO MEFs infected for 2h. (f) and (g) Quantification of the number of Δ*actA L. monocytogenes* positve for ubiquitin (f) and p62 (g) in WT and ULK1 KO MEFs infected for 4h. (h) and (i) Quantification of the number of Δ*actA L. monocytogenes* positive for ubiquitin (h) and p62 (i) in WT and ULK1 KO MEFs infected for 8h. Scale bars are 10μM. Values are mean ± SEM. *p<0.05, two-tailed unpaired t-test. N=3-5.

### Phosphorylation at p62 S409 is required for the recruitment of p62

p62-dependent selective autophagy has been implicated as a compensatory pathway for the degradation of large ubiquitylated protein aggregates. However, the mechanisms that control substrate selection are poorly understood. To date, it is known that post-translational modifications on p62 determine the ability of p62 to promote selective autophagy. For example, it has been shown that casein kinase 2 (CK2)-dependent phosphorylation of p62 at serine 403 (S403) increases the affinity of its UBA domain for polyubiquitylated chains, to promote the removal of ubiquitylated substrates (38). Similarly, TANK-binding kinase 1 (TBK-1) was demonstrated to phosphorylate p62 at the same site, and its pharmacological inhibition of TBK-1 abrogated the autophagic-dependent clearance of *M. tuberculosis* in macrophages (39). Another phosphorylation site on p62 (serine 409, S409) has also been reported to increase the affinity of its UBA domain for ubiquitylated protein aggregates. Interestingly, phosphorylation on S409 was reported to be dependent on ULK1 kinase activity (25). Considering that we demonstrated that ULK1 is necessary for the ubiquitylation of bacteria and p62 recruitment, we argued whether phosphorylation on p62 S409 would also be required to recruit p62 to *L. monocytogenes* surface. For this purpose, we infected MEFs from p62 knockout (p62 KO) mice transduced with a retrovirus for the expression of FLAG-WT p62 or its phosphorylation-null form, in which serine 409 was substituted with alanine (FLAG-p62 S409A). We monitored the recruitment of p62 to the surface of *L. monocytogenes* by immunofluorescence for 2 and 4h and observed a significant reduction in the number of bacteria associated with p62 in MEFs expressing p62 S409A when compared to MEFs expressing WT p62 (**Fig.6A-C**). Quantification of the number of bacteria associated with p62 demonstrated a significant decrease in the recruitment of this protein in cells expressing the phosphorylation-null mutant when compared to MEFs expressing WT p62 (∼17% in WT p62 vs. ∼1% in MEFs expressing p62 S409A **Fig. 6D**). Next, we wondered if the phosphorylation on p62 S409 would also affect the recruitment of p62 by Δ*actA L. monocytogenes*. Immunofluorescence microscopy analysis of MEFs expressing WT p62 and p62 S409A did not reveal differences in the association of p62 to Δ*actA L. monocytogenes* (**Fig. 7A-D**). Overall, these results imply, for the first time, that the phosphorylation on p62 S409A is crucial for efficiently recruiting p62 to the surface of cytosolic ubiquitylated bacteria.

**FIGURE 6:**
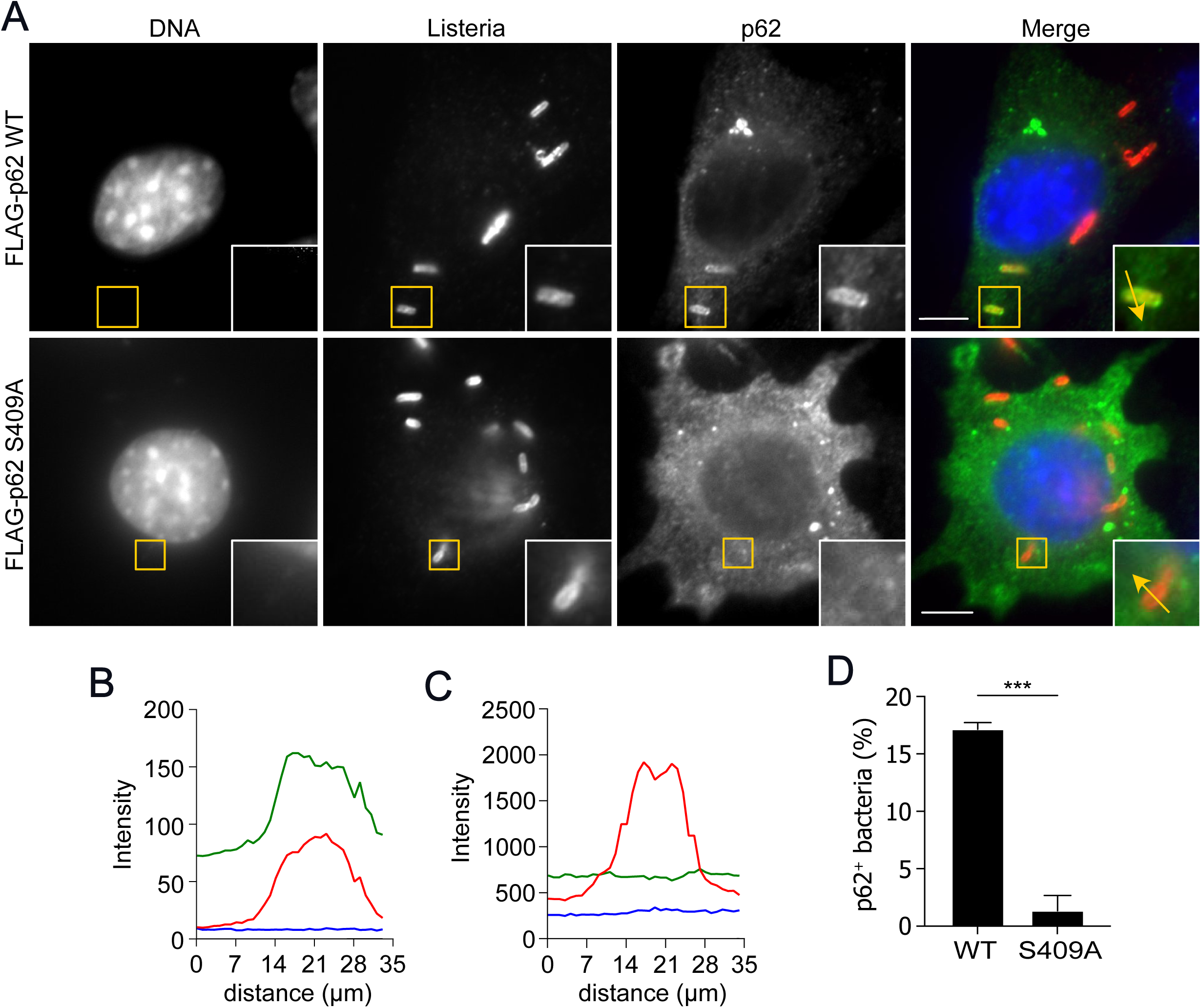
Phosphorylation on p62 S409 in essential for recruitment to *L. monocytogenes*. (a) Representative confocal imagens of p62 KO MEFs transduced with retrovirus for the stable expression of FLAG-p62 WT or FLAG-p62 S409A infected with WT *L. monocytogenes* for 2h and stained for *Listeria* (red), FLAG (green), and DNA (blue). (b) and (c) Plot profiles of fluorescence intensity along the yellow arrow traced in the inset in (a); (b) MEFs p62 KO MEFs expressing FLAG-p62 WT and (c) MEFs p62 KO MEFs expressing FLAG-p62 S409A. FLAG (green line), *Listeria* (red line), and DNA (blue line). (d) Quantification of *L. monocytogenes* positive for p62 in MEFs p62 KO MEFs stably expressing FLAG-p62 WT or FLAG-p62 S409A, infected as described in (a). Scale bars are 10 µM. Values are mean ± SEM. *p<0.05, two-tailed unpaired t-test. N=3

**FIGURE 7:**
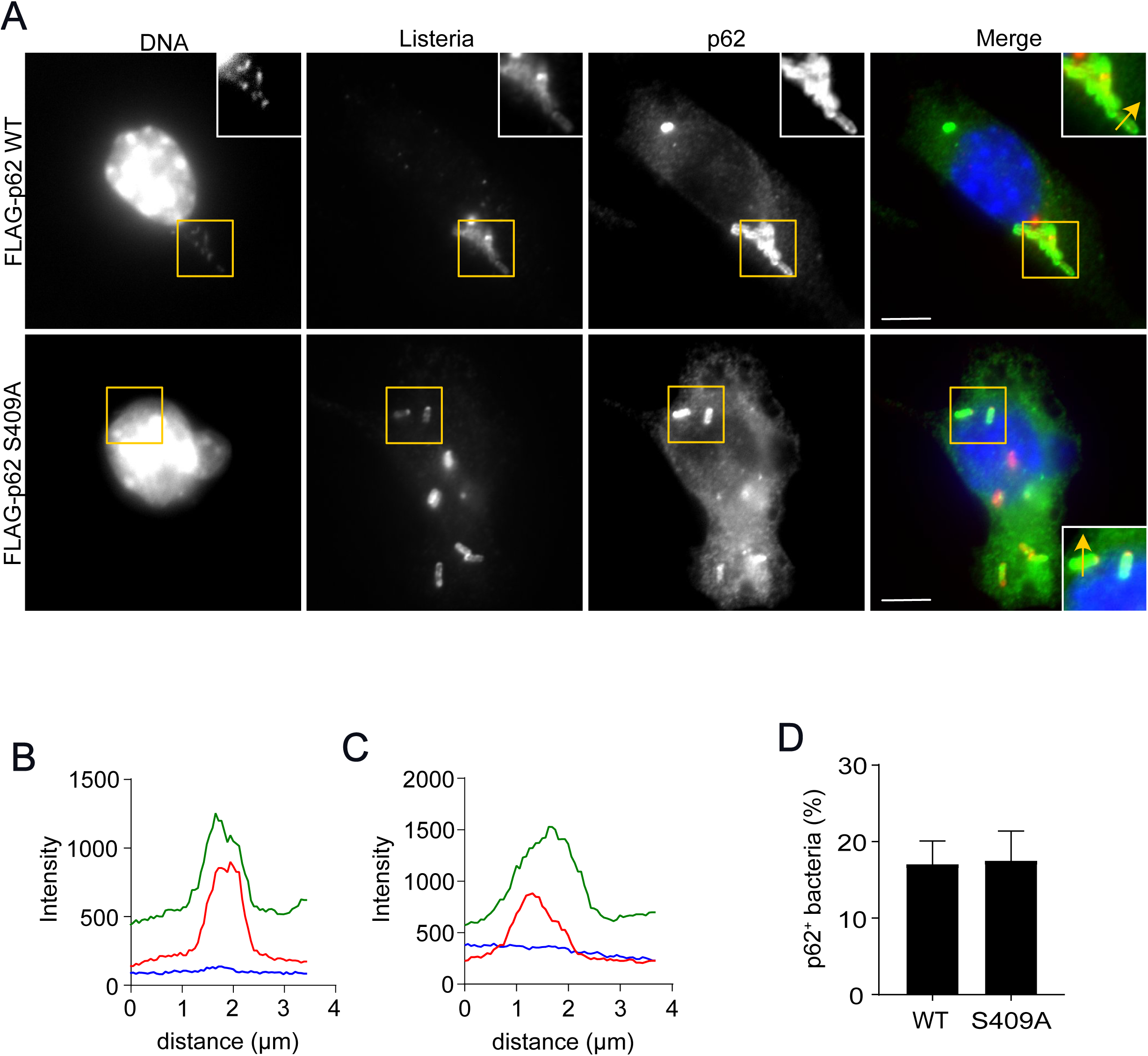
Phosphorylation on p62 S409 is not required to recruit p62 to *L. monocytogenes* lacking actA. (a) Representative confocal images of p62 KO MEFs transduced with retrovirus for the stable expression of FLAG-p62 WT or FLAG-p62 S409A infected with Δ*acta L. monocytogenes* for 2h and stained with antibodies for *Listeria* (red) and FLAG (green), and DNA (blue). Scale bars are 10μM. (b) and (c) Plot profiles of fluorescence intensity along the yellow arrow traced in the inset in (a); (b) MEFs p62 KO MEFs expressing FLAG-p62 WT and (c) MEFs p62 KO MEFs expressing FLAG-p62 S409A. FLAG (green line), *Listeria* (red line), and DNA (blue line). (d) Quantification of Δ*acta L. monocytogenes* positive for p62 in MEFs p62 KO MEFs stably expressing FLAG-p62 WT or FLAG-p62 S409A, infected as described in (a). Values are mean ± SEM. N=3

### ULK1 and phosphorylation at p62 S409 are required for growth restriction of *L. monocytogenes*

During the last decade, xenophagy has emerged as a critical effector of the innate immune system as an essential mechanism against infection of the cytosolic compartment (2). The formation of autophagosomes targeted to cytosolic bacteria has been reported to depend on the addition of ubiquitin and p62 to the surface of cytosolic bacteria. Our present data demonstrate that ULK1 is a major player in these early steps of xenophagy by controlling Ub and p62 tagging of *L. monocytogenes*. As such, we questioned whether the lack of ULK1 would affect the restriction of intracellular replication of the bacteria. To test this, we quantified the number of cytosolic bacteria in WT and ULK1 KO MEFs infected with *L. monocytogenes*. As demonstrated in **Figure 8A** and **B**, at 8h post-infection, ULK1 KO MEFs harbored significantly higher numbers of cytosolic bacteria as compared to WT MEFs (∼113 bacteria per cell in ULK1 KO MEFs vs. ∼31 bacteria per cell in WT MEFs). Considering the results described above, we aimed to investigate if phosphorylation of p62 at S409 was also implicated in the intracellular restriction of *L. monocytogenes* replication. Immunofluorescence microscopy of p62 KO MEFs expressing WT p62 or p62 S409A infected with *L. monocytogenes* for the quantification of the number of bacteria present in the cytosol showed significantly higher numbers of bacteria in MEFs expressing p62 S409A (∼26 bacteria per cell in p62 S409A expressing MEFs vs. 16 bacteria per cell in WT p62 expressing MEFs, **Fig. 8C** and **D**). Together, these results demonstrate that ULK1 and phosphorylation of p62 S409 are essential for efficiently restricting bacterial replication in the cytosol.

**FIGURE 8:**
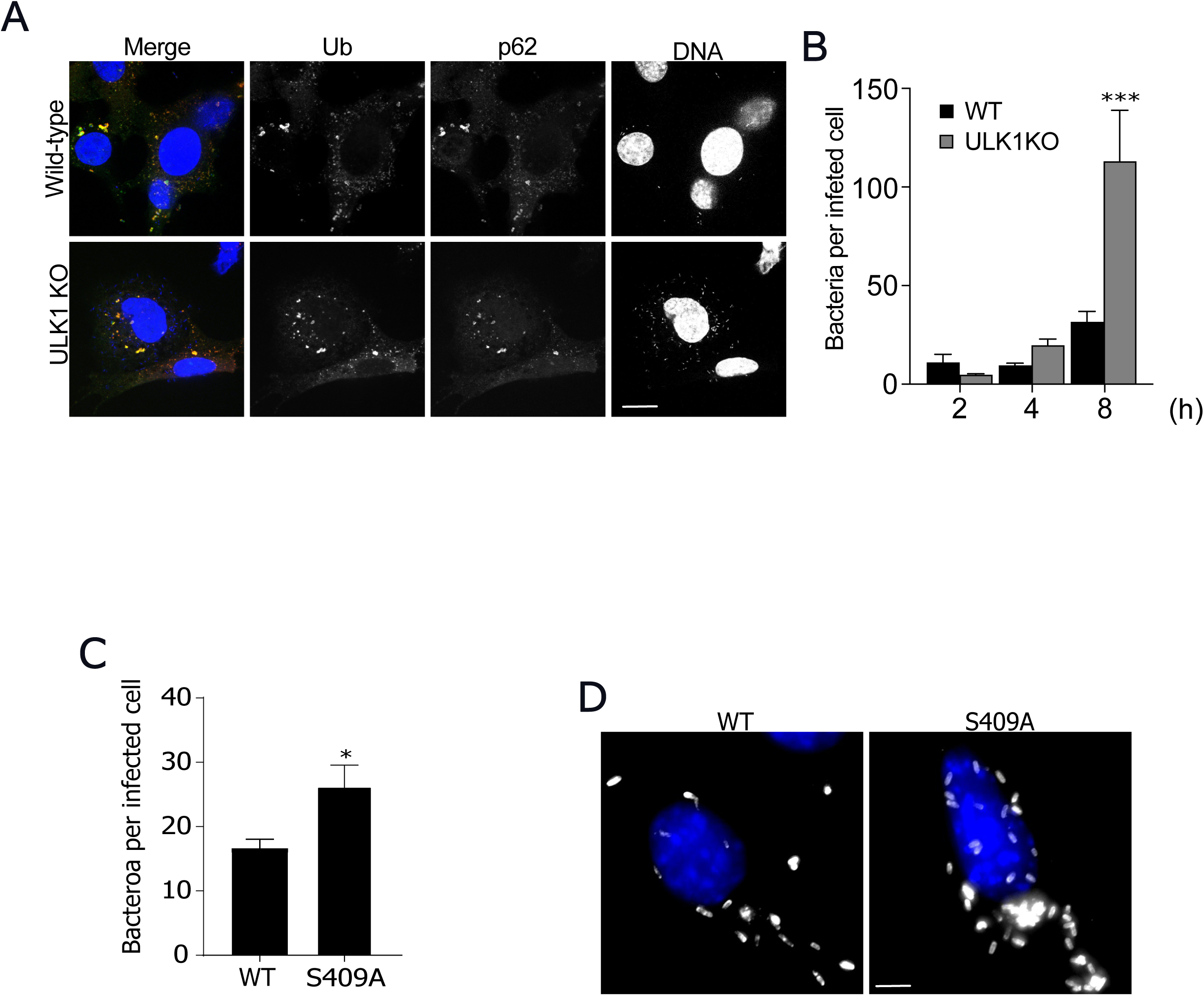
ULK1 and phosphorylation at p62 S409 afford the control of intracellular replication of *L. monocytogenes*. (a) Representative confocal images of WT and ULK1 KO MEFs infected with *L. monocytogenes* for 8h and stained for ubiquitin (green), p62 (red), and DNA (blue). Scale bars are 1μM. (b) Quantification of the number of intracellular bacteria in WT and ULK1 KO MEFs infected with *L. monocytogenes* for 2, 4, 8h. Values are mean ± SEM. ***p<0.0001, one-way ANOVA with Sidak’s post-test. N=2. (c) Quantification of the number of intracellular bacteria in p62 KO MEFs stably expressing FLAG-p62 WT or FLAG-p62 S409A infected with *L. monocytogenes* for 4h. Values are mean ± SEM. *p<0.05, two-tailed unpaired t-test. (d) Representative confocal images of p62 KO MEFs stably expressing FLAG-p62 WT or FLAG-p62 S409A infected with *L. monocytogenes* for 4h and stained for DNA (blue) and *Listeria* (gray). Scale bars are 5μM. N=3

## DISCUSSION

Xenophagy has emerged as an essential mechanism for the control of cytosolic bacteria. In this context, despite major advances in the study of the interactions between autophagy machinery and cytosolic bacteria, the mechanisms underlying the recognition of bacteria by autophagic adapters are still largely undefined. Although several studies demonstrate the importance of recruiting Ub and p62 to bacterial surfaces, the factors regulating these crucial steps of xenophagy are far from elucidated. Here, we used *L. monocytogenes* as a bacterial model to exploit factors driving the recruitment of Ub and p62 during infection of non-myeloid cells. Our results demonstrate that a significant proportion of the cytosolic *L. monocytogenes* population is tagged by autophagy proteins such as LC3, Ub, and p62 in a LLO-dependent manner, in agreement with data from the literature (24). To understand the factors that control the ubiquitylation and p62 recruitment, we sought to investigate the role of ULK1, a major component of the ULK1 complex, known to integrate signals from energy sensing pathways that have been previously reported to be involved in autophagy triggering during *L. monocytogenes* infection (32). Currently, the knowledge on the role of ULK1 in the autophagic response to intracellular bacteria is limited to its participation in the generation of a replicative niche for *Staphylococcus aureus* (40, 41) or *Salmonella* (42) by initiating autophagosome membrane formation. Here, we expand the role ULK1, demonstrating that its deficiencyleads to a severe impairment of bacterial ubiquitylation during xenophagy of *L. monocytogenes*. Our results expand the role of ULK1 in xenophagy beyond autophagosome formation, implying its participation in early steps of the process, such as ubiquitylation of bacterial surfaces. Based on previous reports demonstrating that the ULK1 complex is recruited to the bacterial surface of *Salmonella* (42), it is tempting to speculate that ULK1 might also be present on *Listeria* surface to coordinate the recruitment of the ubiquitin-ligases such as Parkin and SMURF1, previously implicated in the ubiquitylation of cytosolic bacteria (10, 11).

Early reports demonstrated that ULK1 controls the phosphorylation of p62 at S409 during proteotoxic stress (25). According to the authors, ULK1-dependent phosphorylation of p62 increases the affinity of its UBA domain for ubiquitin. In light of our results demonstrating that ULK1-deficient cells are limited in their ability to efficiently promote the ubiquitylation and p62 recruitment to cytosolic *Listeria*, we decided to investigate if phosphorylation at p62 S409 would be essential for the recognition of ubiquitin on the surface of *L. monocytogenes*. Our results show that cells harboring a p62 phosphorylation null mutant in this residue display a significantly reduced capacity to localize with the wild-type strain. The data presented here raises the possibility that ULK1-dependent control of xenophagy occurs at two different moments, i.e., in the regulation of ubiquitylation step and p62 S409 phosphorylation. However, we must note that we cannot rule out the possibility that the reduced recruitment of p62 observed in ULK1-deficient MEFs was not a mere consequence of the diminished ubiquitylation. Despite a previous report describing the TBK1-dependent phosphorylation of p62 S403 during *M. tuberculosis* infection (38), our results are the first demonstration that post-translational modifications on p62 affect the recognition of ubiquitylated cytosolic bacteria.

It is interesting to note that while ULK1-deficiency and blockade of p62 S409 phosphorylation deeply affected host cell ability to recruit Ub and p62 during the infection with the wild-type strain, it did not impact the presence of these two proteins on the surface of Δ*actA L. monocytogenes*. Although the ActA-dependent mechanism involved in the escape from autophagy remains to be fully clarified, our results raise the possibility that ActA expression might contribute to avoiding the localization of ULK1 and a putative ULK1-dependent recruitment of ubiquitin-ligases to the vicinity of *L. monocytogenes*, blocking the ULK1-dependent ubiquitylation and p62 S409 phosphorylation.

Finally, our experiments demonstrating that both ULK1-deficiency and blockade of p62 S409 phosphorylation lead to impaired control of bacterial replication, together with previous work showing that the small-molecule autophagy inducer A77 1726 activates the AMPK-ULK1 axis to restrict the intracellular growth of *Salmonella* (42) reinforce the importance of our data and highlight the potential of ULK1 as a target for future therapeutic approaches to fight infection with intracellular bacteria.

## EXPERIMENTAL PROCEDURES

### Antibodies and reagents

anti-p62 (ab109012) and anti-*Listeria monocytogenes* (ab35132) were from Abcam (Cambridge, UK). Anti-ubiquitin (ADI-SPA-203-F) was from Enzo Life Sciences (Farmingdale, NY, USA). anti-LAMP1 (sc-20011) and anti-mTOR1 (2983) were from Santa Cruz Biotechnology (Dallas, TX, USA) and Cell Signalling (Danvers, MA, USA). Alexa Fluor™ 488 goat anti-mouse IgG, Alexa Fluor™ 488 goat anti-rat IgG, and Alexa Fluor™ 568 goat anti-rabbit IgG were from Invitrogen. Anti-FLAG, Puromycin, and DAPI were from Sigma-Aldrich (Saint Louis, MO, USA). Prolong Gold Antifade reagent was from Invitrogen.

### Bacterial strains and cell culture

wild-type *L. monocytogenes* 10403S, Δ*hly L. monocytogenes* 10403S, and Δ*actA L. monocytogenes* 10403S were kindly donated by Dr. Daniel Portnoy and Dr. Gabriel Mitchell. (Department of Molecular and Cell Biology, University of California, Berkeley, CA, USA). All bacterial strains were grown in brain-heart infusion (BHI) broth from Kasvi (Pinhais, Brazil). p62 KO MEFs retrovirally transduced with FLAG-p62 WT or FLAG-p62 S409A were a gift from Dr. Zhenyu Yue (Department of Neurology and Neuroscience, Friedman Brain Institute, Icahn School of Medicine at Mount Sinai, New York, NY, USA). Wild-type and ULK1 KO MEFs were donated by Dr. Douglas Green (Department of Immunology Comprehensive Cance Center St. Jude Graduate School of Biomedical Sciences). Expression of GFP-LC3 in MEFs was performed as previously described (Travassos 2010). All cells were cultured in Dulbecco’s modified Eagle medium (DMEM) supplemented with 10% fetal calf serum (FCS, Invitrogen, Waltham, MA, USA), 2 mM L-glutamine, 50 IU penicillin/50 mg/ml streptomycin (Invitrogen), and plasmocin (Invivogen, Toulouse France). p62 KO MEFs transduced with retrovirus for FLAG-p62 WT and FLAG-p62 S409A were kept in puromycin (3 μg/ml). Cells were maintained in 95% air 5% CO_2_ at 37°C.

### Bacterial infections

MEFs were seeded in 24-well plates containing glass coverslips at a density of 1 x 10^5^ cells/well in antibiotic-free medium. *L. monocytogenes* strains grown to exponential phase were centrifuged at 2000 *g* and adjusted to to infect MEFs at a MOI of 100 for 1h in 95% air, 5% CO 2 at 37°C. After this period cells were washed twice with FSC-free medium and treated with gentamicin-containing complete medium (50 μg/ml) for the indicated time points.

### Immunofluorescence microscopy

after infection, with 4% (wt/vol) paraformaldehyde for 15 minutes at room temperature, blocked with PBS-BSA (1% wt/vol), permeabilized with Triton X-100 (Sigma-Aldrich), and incubated with primary antibodies for 2h. Cells were then washed three times with PBS and stained with the appropriate secondary antibodies and DAPI.

### Image analysis

association of GFP-LC3, Ub, and p62 with *L. monocytogenes* was counted from confocal images acquired with a Cell Observer Yokogawa spinning disk microscope (Zeiss, Rostock, Germany). The number of associations was quantified using Image J software (Schneider 2012). Plot profiles were generated using the Zen Desk software from (Zeiss).

### Statistical analysis

Results are expressed as mean ± SEM of data obtained in independent experiments. Statistical differences between groups were determined using Student’s *t*-test. Statistical significance was set at *P* < 0.05.

## Funding

LHT lab was supported by grants from Fundação Carlos Chagas de Amparo à Pesquisa do Estado do Rio de Janeiro (FAPERJ, Programa Cientista do Nosso Estado (CNE), #E26/200.998/2022 and Programa de Auxílio Básico à Pesquisa (APQ1) #E26/211.466/2021) and Conselho Nacional de Desenvolvimento Científico e Tecnológico (CNPq). LAMC lab was supported by grants from FAPERJ Programa Cientista do Nosso Estado (CNE), (#E-26/200.953/2021) and Conselho Nacional de Desenvolvimento Científico e Tecnológico (CNPq). MSS and JSR were supported by fellowships from Coordenação de Aperfeiçoamento de Pessoal de Nível Superior (CAPES)/Biocomputacional, TSMF, and RMR were supported by scholarships from CNPQ. TDP was supported by a Programa de Bolsas Inciação Científica – UFRJ studentship. RMCS was supported by post-doctoral fellowships from CAPES/Epidemias and a grant from Fundação Carlos Chagas de Amparo à Pesquisa do Estado do Rio de Janeiro (FAPERJ, Programa Jovem Pesquisador (E-26/200.628/2022.)

---

**SUPPLEMENTARY FIGURE 1:**
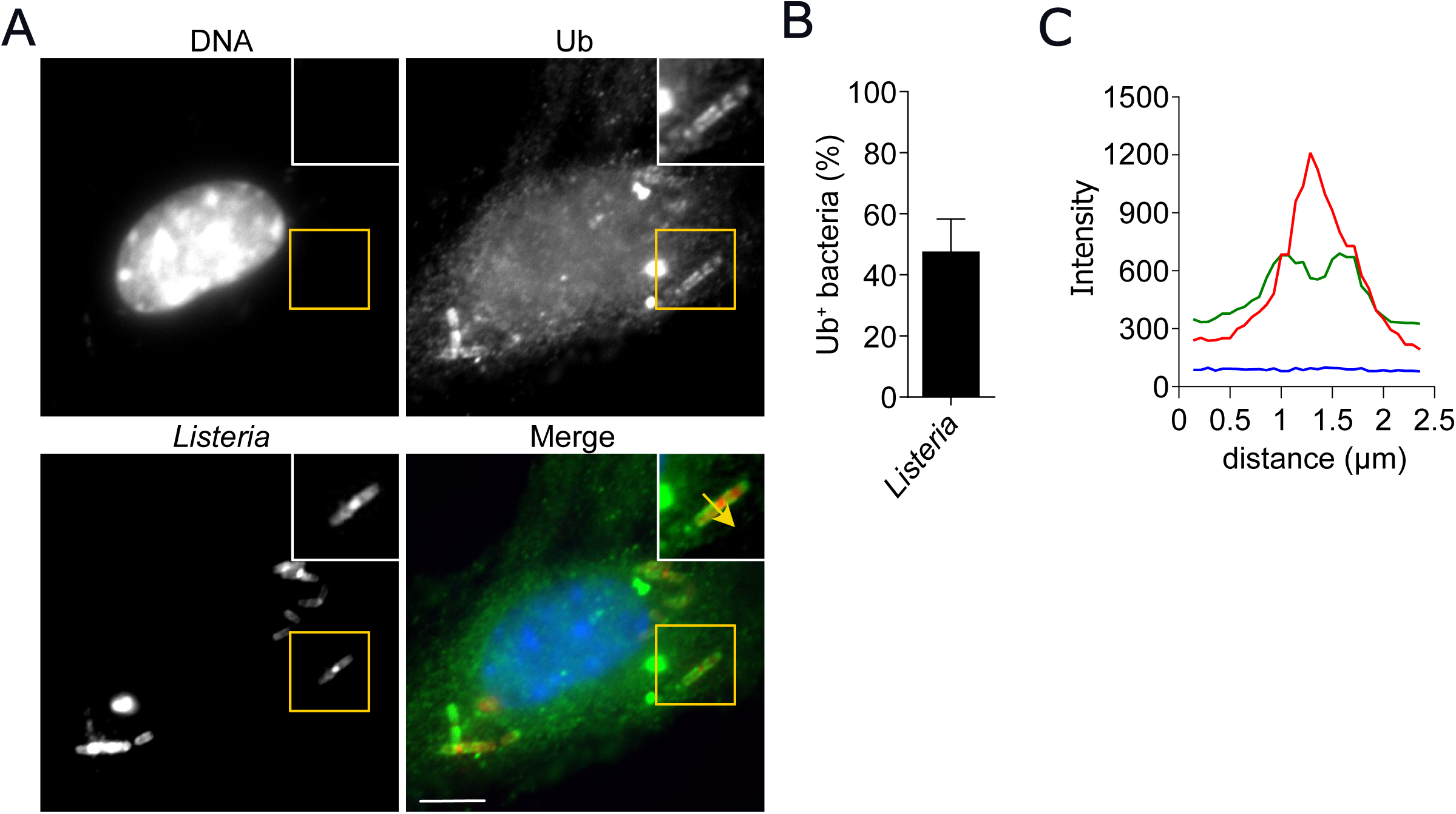
Intracellular *L. monocytogenes* is ubiquitylated. (a) Representative confocal images of MEFs infected with WT *L. monocytogenes* for 2h and stained for ubiquitin (green), *Listeria* (red), and DNA (blue), N=2. (b) Quantification of the number of *L. monocytogenes* positive for ubiquitin in experiments realized as in (a), N=2. (c) Plot-profiles of fluorescence intensity along the yellow arrow traced in the inset in (a). Values are means ± SEM. Scale bars are 10μM

**SUPPLEMENTARY FIGURE 2:**
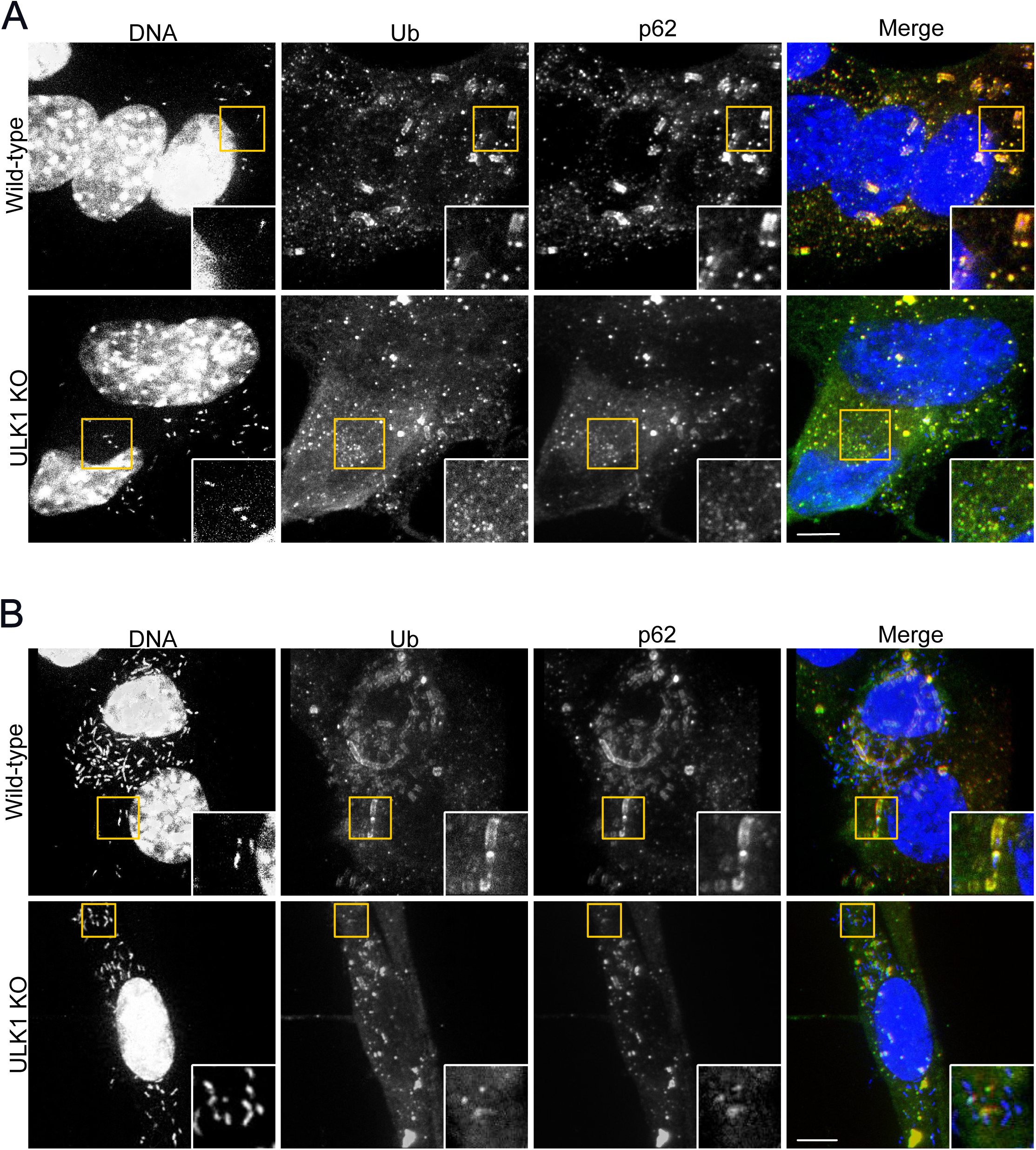
ULK1 is required for the efficient recruitment of ubiquitin and p62 to *L. monocytogenes* surface. (a) and (b) Representative confocal images of WT and ULK1 KO MEFs infected with WT *L. monocytogenes* for 4h (a) and 8h (b) and stained for ubiquitin (green), p62 (red), and DNA (blue). N=5. Scale bars are 10μM.

**SUPPLEMENTARY FIGURE 3:**
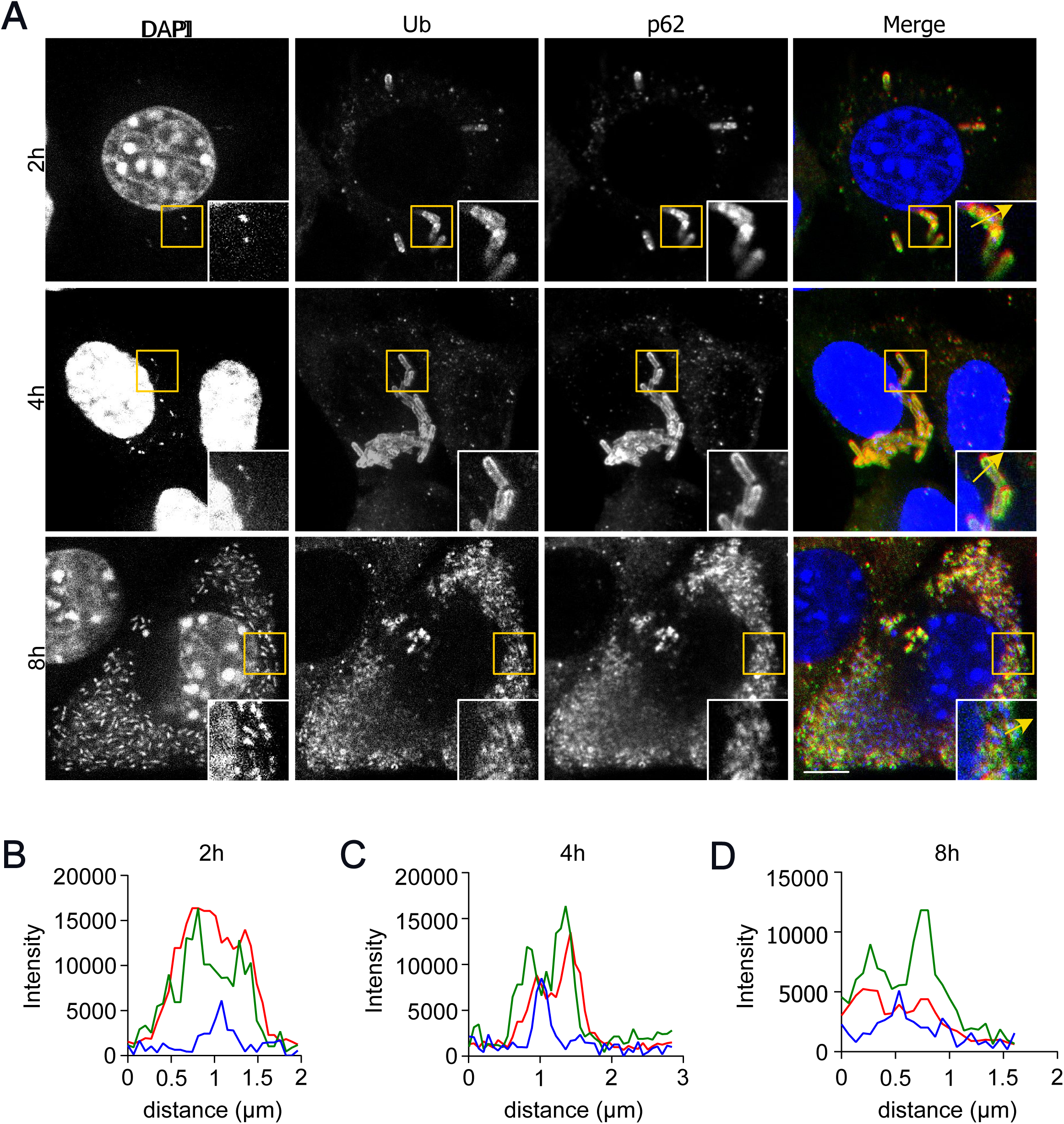
*ΔactA* mutants are efficiently ubiquitylated and targeted by p62. (a) Representative confocal images of WT MEFs infected with Δ*actA L. monocytogenes* mutant for 2h (top), 4h (center), and 8h (bottom) and stained for ubiquitin (green), p62 (red), and DNA (blue). (b), (c), and (d) Plot profiles of fluorescence intensity along the yellow arrows traced in the insets in (a) in WT MEFs infected with *ΔactA L. monocytogenes* for 2h (a), 4h (b), and 8h (c). Scale bars are 10μM.

**SUPPLEMENTARY FIGURE 4:**
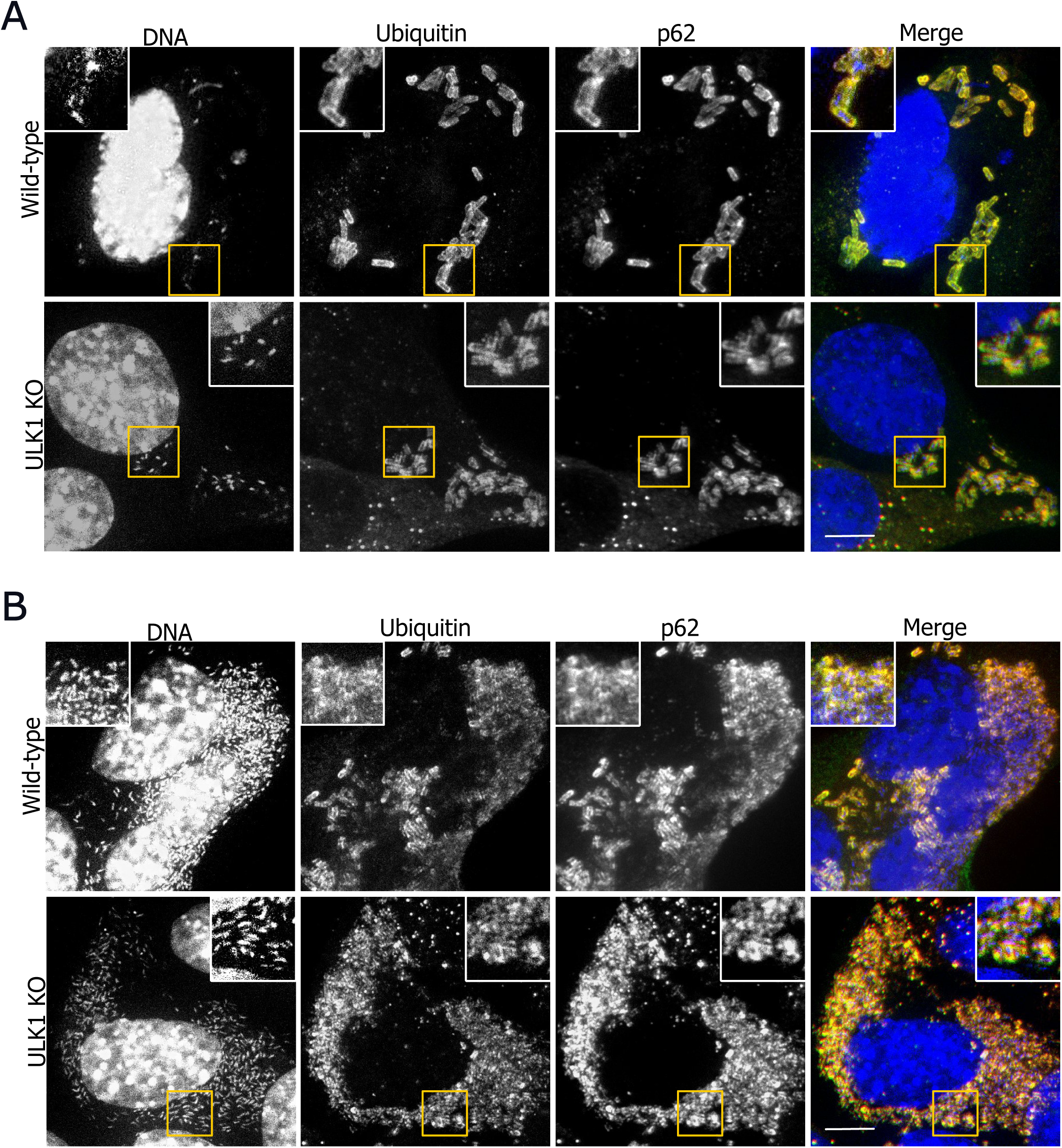
Ubiquitin and p62 recruitment to Δ*actA L. monocytogenes* surface do not require ULK1. (a) and (b) Representative confocal images of WT and ULK1 KO MEFs infected with Δ*actA L. monocytogenes* for 4h (a) and 8h (b) and stained for ubiquitin (green), p62 (red), and DNA (blue).. Scale bars are 10μM. N=5

